# Combination of tenofovir and emtricitabine with efavirenz does not moderate inhibitory effect of efavirenz on mitochondrial function and cholesterol biosynthesis in human T lymphoblastoid cell line

**DOI:** 10.1101/297044

**Authors:** Min Li, Anuoluwapo Sopeyin, Elijah Paintsil

## Abstract

Efavirenz (EFV), the most popular non-nucleoside reverse transcriptase inhibitor, has been associated with mitochondrial dysfunction in most in vitro studies. However, in real life the prevalence of EFV-induced mitochondrial toxicity is relatively low. We hypothesized that the agents given in combination with EFV might moderate the effect of EFV on mitochondrial function. To test this hypothesis, we cultured human T lymphoblastoid cell line (CEM cells) with EFV alone and in combination with emtricitabine (FTC) and tenofovir disoproxil fumarate (TDF) to investigate the effects on mitochondrial respiration and function and cholesterol biosynthesis.

There was a statistically significant concentration- and time-dependent apoptosis, reduction in mitochondrial membrane potential (ΔΨ), and increase production of reactive oxygen species (ROS) in cells treated with either EVF alone or in combination with TDF/FTC. EFV treated cells compared to DMSO treated cells had significant reduction in oxygen consumption rate (OCR) contributed by mitochondrial respiration, ATP production-linked respiration, and spare respiratory capacity (SRC). Treatment with EFV resulted in a decrease in mitochondrial DNA content, and perturbation of more coding genes (n=13); among these were 11 genes associated with lipid or cholesterol biosynthesis. Our findings support the growing body of knowledge on the effects of EFV on mitochondrial respiration and function and cholesterol biosynthesis.

Interestingly, combining TDF/FTC with EFV did not alter the effects of EFV on mitochondrial respiration and function and cholesterol biosynthesis. The gap between the prevalence of EFV-induced mitochondrial toxicity *in vitro* and *in vivo* studies may be explained by individual differences in the pharmacokinetic of EFV.

## INTRODUCTION

Combination antiretroviral therapy (ART) has resulted in a decrease in AIDS-related morbidity and mortality (1), though the therapeutic benefit of ART is often limited by delayed toxicity (2). Although contemporary ART regimens compared to early ART regimens are less toxic(3, 4), toxicity is still pervasive and affects a significant number of people living with HIV (4, 5). *In vitro* studies demonstrated that inhibition of polymerase gamma (Pol-γ), enzyme responsible for mitochondrial DNA replication, by nucleoside reverse transcriptase inhibitors (NRTIs) leads to depletion of mitochondrial DNA (mtDNA) and subsequent mitochondrial dysfunction(6, 7); the “Pol-γ inhibition” hypothesis. However, there is a growing body of knowledge to suggest that ART-associated mitochondrial dysfunction cannot be explained solely by Pol-γ inhibition(8, 9). For instance, other classes of ART such as protease inhibitors (PIs) and non-nucleoside reverse transcriptase inhibitors (NNRTIs) do not inhibit Pol-γ and yet they cause side effects akin to mitochondrial dysfunction(8, 10). Taken together, there must be alternative or additional mechanisms by which ART impairs mitochondrial function.

Efavirenz (EFV), the most popular NNRTI and a key component of several ART regimens, has been associated with metabolic disorders(11), hepatic toxicity (12, 13), diminished bone density (14), neuropsychiatric symptoms(15, 16), and neurocognitive impairment (17).

Although the underlying cellular and molecular mechanisms of EFV-induced toxicity are still not well understood, several *in vitro* and *in vivo* studies have implicated mitochondrial dysfunction as the underlying mechanism. EFV affections on mitochondria include decrease in mitochondrial membrane potential, inhibition of OXPHOS complex I enzymes, decrease in oxygen consumption, and increase production of mitochondrial reactive oxygen species (ROS)(8, 18, 19).

With this litany of effects of EFV on mitochondria, one would have expected a much higher incidence of EFV-associated toxicity in patients. The incidence of severe EFV-associated neuropsychiatric symptoms is less than 2% of patients (15, 20); and severe hepatic toxicity is up to 8% of patients (12, 13). In real life, EFV is given in combination with other antiretroviral agents. We, therefore, hypothesized that the agents given in combination with EFV might moderate the effect of EFV on mitochondrial function; hence, the relatively low incidence of EFV-induced mitochondrial toxicity in patients. To test this hypothesis, we cultured human T lymphoblastoid cell line (CEM cells) with EFV alone and in combination with emtricitabine (FTC) and tenofovir disoproxil fumarate (TDF) to investigate the effects on mitochondrial function and cholesterol biosynthesis.

## RESULTS

### EFV treatment decreased CEM cell growth

We treated CEM cells with EFV, TDF/FTC, or TDF/FTC/EFV at multiples of their respective C_max_ (i.e., 1x-, or 2x-C_max_). Growth of cells treated with antiretroviral (ARV) drugs compared to DMSO-treated cells was monitored. All the ARVs tested affected CEM cell growth to a certain degree in a concentration- and time-dependent manner except for TDF/FTC combination (Figure 1A). At day 1, EFV treatment (1x-, and 2x-C_max_) reduced cell growth more than TDF/FTC/EFV combination treatment. At day 2, the effect of treatment with EFV alone on cell growth was comparable to that of treatment with TDF/FTC/EFV combination.

**Figure 1.**
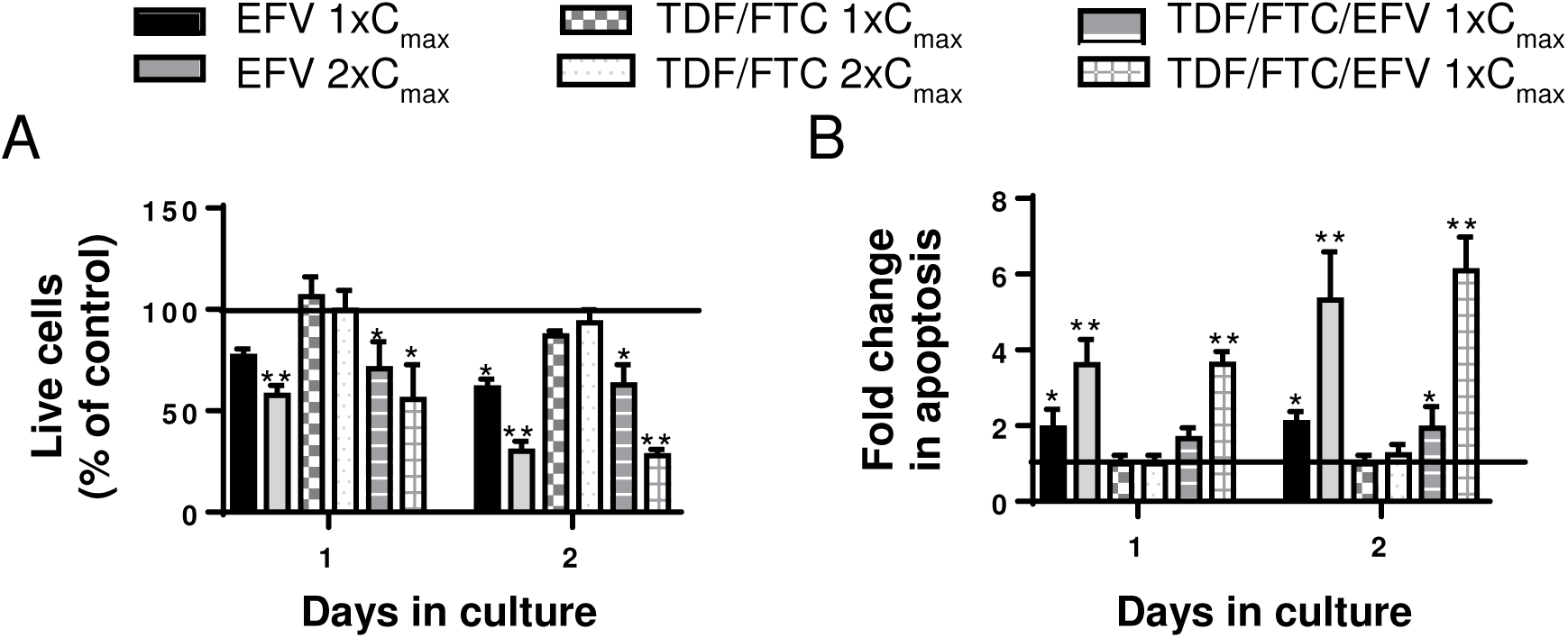
Effect of efavirenz on cell viability and apoptosis in of CEM cells Human T lymphoblastoid cell line (CEM cells) was treated with multiplicities of the plasma peak concentrations (1x-, or 2xC_max_) of Efaverinz (EFV), Tenofovir and Emtricitabine (TDF/FTC), and TDF/FTC/EFV. A. Each day, aliquots of the cell culture were collected and live cells were counted with Trypan blue staining using hemocytometer, percentage of live cells was compared with DMSO-treated control cells. B. Apoptosis was assessed by staining with an FITC Annexin V Apoptosis Detection Kit I. Bar graphs represent proportion of apoptotic cells after 1 and 2 days of treatment compared to DMSO-treated cells. Data represent at least 3 independent experiments and plotted as mean ± SD. P-values are two sided and considered significant if <0.05 (*), < 0.01 (**), or < 0.001 (***>).

### EFV treatment increased proportion of apoptotic cells

Mitochondria are central to the process of cell apoptosis. We, therefore, investigated the effect of exposure of CEM cells to EFV, TDF/FTC, or TDF/FTC/EFV on apoptosis. We determined cell death/apoptosis using PI/Annexin-V flow cytometry at day 1 and day 2 (21). Figure 1B illustrates the fold-change in apoptosis in ARV treated cells compared to DMSO treated cells. We observed a statistically significant concentration- and time-dependent apoptosis in cells treated with either EVF alone or in combination with TDF/FTC. There was no statistically significant difference between cells treated with TDF/FTC at the two concentration and cells treated with DMSO.

### EFV treatment decreased mitochondrial membrane potential in cell culture

We next investigated the effect of EFV, TDF/FTC, or TDF/FTC/EFV treatment on mitochondrial membrane potential (ΔΨ) of CEM cells using TMRE (tetramethylrhodamine, ethyl ester). Compared to cells treated with DMSO, cells treated with both concentrations of EFV or TDF/FTC/EFV showed statistically significant higher proportion of cells with decreased mitochondrial ΔΨ after 1 and 2 days in culture in a dose- and time-dependent manner (Figure 2A). We did not observe any statistically significant difference between cells treated with TDF/FTC at the two concentrations compared to cells treated with DMSO.

**Figure 2.**
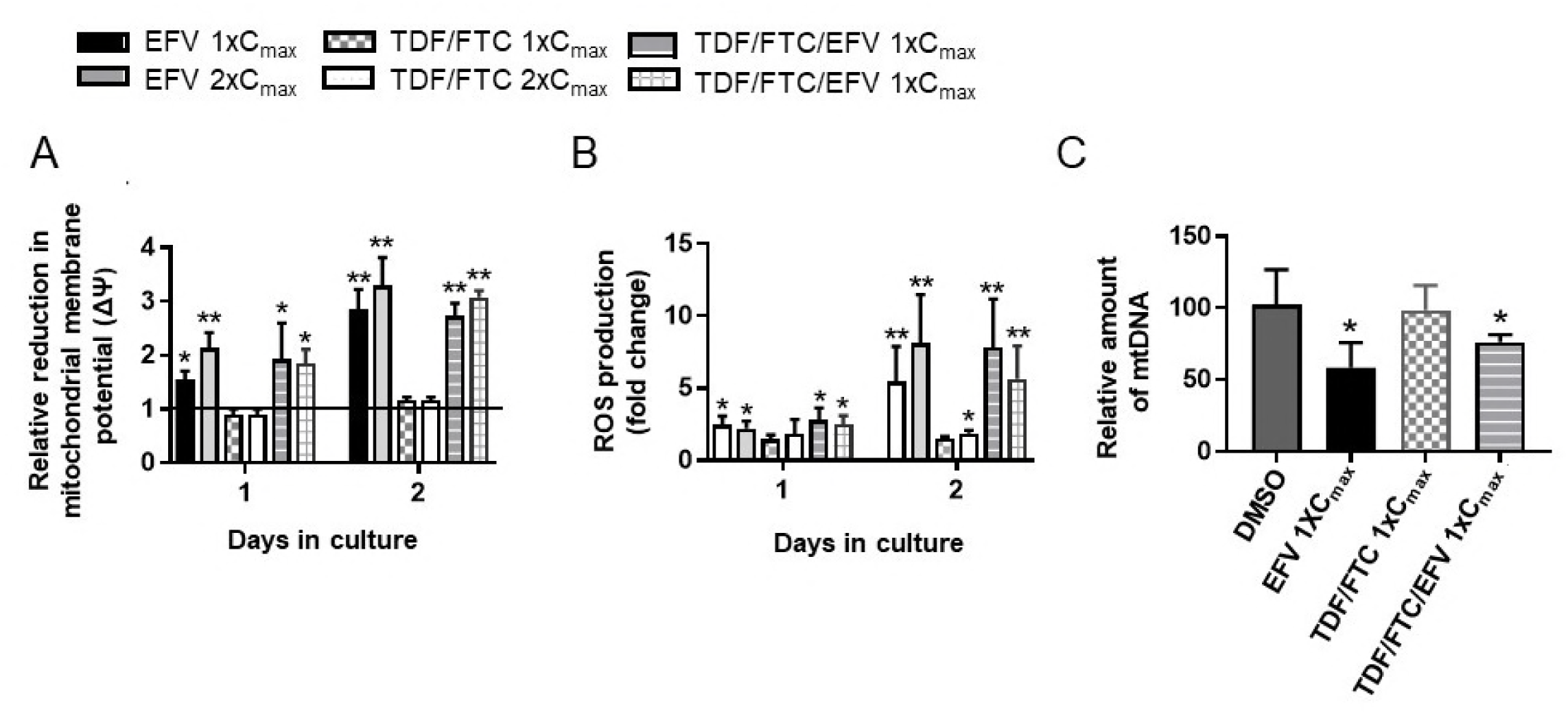
Effect of efavirenz treatment on mitochondrial membrane potential, ROS production and mitochondrial DNA content. Human T lymphoblastoid cell line (CEM cells) was treated with multiplicities of the plasma peak concentrations (1x-, or 2xC_max_) of Efavirenz (EFV), Tenofovir and Emtricitabine (TDF/FTC), and TDF/FTC/EFV. A. At day 1 and 2 the cells were harvested and interrogated for mitochondrial membrane potential (ΔΨ) using TMRE-Mitochondrial Membrane Potential Assay Kit with Flow cytometry. Inactive mitochondria had decreased membrane potential and failed to sequester TMRE. The bar graph represents the relative decrease of mitochondrial ΔΨ in cells treated with antiretroviral drugs compared to DMSO-treated control cells. B. Production of ROS was determined using MitoSOX™ assay. The cells were washed once with wash buffer before incubation with 5 μM MitoSOX™ reagent working solution at room temperature for 10 min. The cells were then washed once before running on LSRII flow cytometry (Beckman Coulter). The bar graph represents the fold change in ROS production in cells treated with antiretroviral drugs compared to DMSO-treated control cells. C. The relative mtDNA content was determined on day 2 of culture as the ratio of mtDNA to nuclear DNA using quantitative PCR. Data represent at least 3 independent experiments and plotted as mean ± SD. P-values are two sided and considered significant if <0.05 (*), < 0.01 (**), or < 0.001 (***).

### EFV treatment increased mitochondrial ROS production

Loss of mitochondrial ΔΨ is associated with oxidative stress and, therefore, increased mitochondrial production of ROS(22). We next investigated whether the decreased in mitochondrial ΔΨ in cells treated with EFV translated to an increase in production of ROS. ROS production was determined using MitoSOX™ Red mitochondrial superoxide indicator. As illustrated in Figure 2B, cells treated with EFV and TDF/FTC/EFV produced significantly higher ROS compared to cells treated with DMSO in a time-dependent manner. There was no statistically significant production of ROS in cells treated with TDF/FTC combination on day 1.

### EFV treatment reduced mitochondrial DNA (mtDNA) content

With the effect of EFV treatment alone and in combination with TDF/FTC on cell growth, apoptosis, mitochondrial ΔΨ, and ROS production presented above, we investigated whether these findings are associated with changes in mtDNA content. We determined the mtDNA content on day 2 of CEM cells treated with EFV, TDF/FTC, or TDF/FTC/EFV compared to cells treated with DMSO (Figure 2C). We observed a statistically significant decrease in the mtDNA of CEM cells treated with 1x-C_max_ of EFV and TDF/FTC/EFV. There was no significant change in mtDNA in cells treated with TDF/FTC compared to cells treated with DMSO.

### EFV altered mitochondrial respiratory function of CEM cells

We next used a more sensitive extracellular flux analysis to investigate the effect of EFV on mitochondrial respiratory parameters (23) –basal respiration rate, ATP production-linked rate, proton leakage rate, maximum respiratory capacity (MRC), spare respiratory capacity (SRC), and respiratory control ratio (RCR)—in CEM cells. Sequential addition of inhibitors of the electron transport chain (ETC) –oligomycin, carbonyl cyanide-4-(trifluoromethoxy)phenylhydrazone, FCCP, and a mixture of rotenone and antimycin—was used to obtain data on the relative oxygen consumption rate (OCR) of different components of mitochondrial respiratory function. The relative OCR was computed using the area under the curve of each component of mitochondrial function as illustrated in Figure 3A. Treatment with EFV at the two concentrations decreased mitochondrial respiration by about 22% (Figure 3B). Treatment with TDF/FTC at 1xC_max_ had no effect on mitochondrial respiration, however, treatment with 2xC_max_ led to a 25% increase in mitochondrial respiration. EFV at 1xC_max_ and 2xC_max_ concentrations decreased the ATP production-linked respiration by 51% and 39%, respectively. However, there was no significant effect of any of the combination treatment on ATP-production-linked respiration. Treatment with TDF/FTC/EFV at 2xC_max_ reduced MRC by 27%%. All the treatment conditions resulted in significant decrease in SRC (ranging from 23% to 44%) compared to DMSO treatment, except treatment with 1xC_max_ of TDF/FTC. We computed the parameters in comparison to basal respiration, expressed as a percentage of basal respiration (Figure 3C), where basal respiration is the combination of mitochondrial and non-mitochondrial respirations. In DMSO treated cells, about 70% of the basal respiration was used for ATP production, however, this percentage dropped to 50% with EFV treatment at the two concentrations (Figure 3C). TDF/FTC treatment did not have an effect on the percentage of basal respiration used to ATP production. Only treatment with 2xC_max_ TDF/FTC and 2xC_max_ TDF/FTC/EFV had significant decrease in the percentage of MRC used for basal respiration (p<0.05). Treatment with 1xC_max_ EFV and 2xC_max_ TDF/FTC/EFV caused a significant increase in proportion of proton leak contributing to basal respiration. All treatments tended to reduce the relative contribution of SRC to basal respiration, however, only treatment with 1xCmax TDF/FTC, 2xC_max_ TDF/FTC, and 2xC_max_ TDF/FTC/EFV led to statistically significant reduction (p<0.05). Treatment with 1xC_max_ and 2xC_max_ EFV increased the contribution of non-mitochondrial respiration to basal respiration. The RCR is a sensitive indicator of mitochondrial dysfunction that is influenced by all components of ETC (18). RCR was computed as the ratio between the maximal respiration and the respiration rate detected in the presence of oligomycin (i.e., ATP production-linked respiration). All the treatment conditions tended to decrease RCR compared to DMSO treatment, however, only treatment with 2xC_max_ of TDF/FTC/EFV resulted in a statistically significant decrease in RCR (Figure 3D).

**Figure 3.**
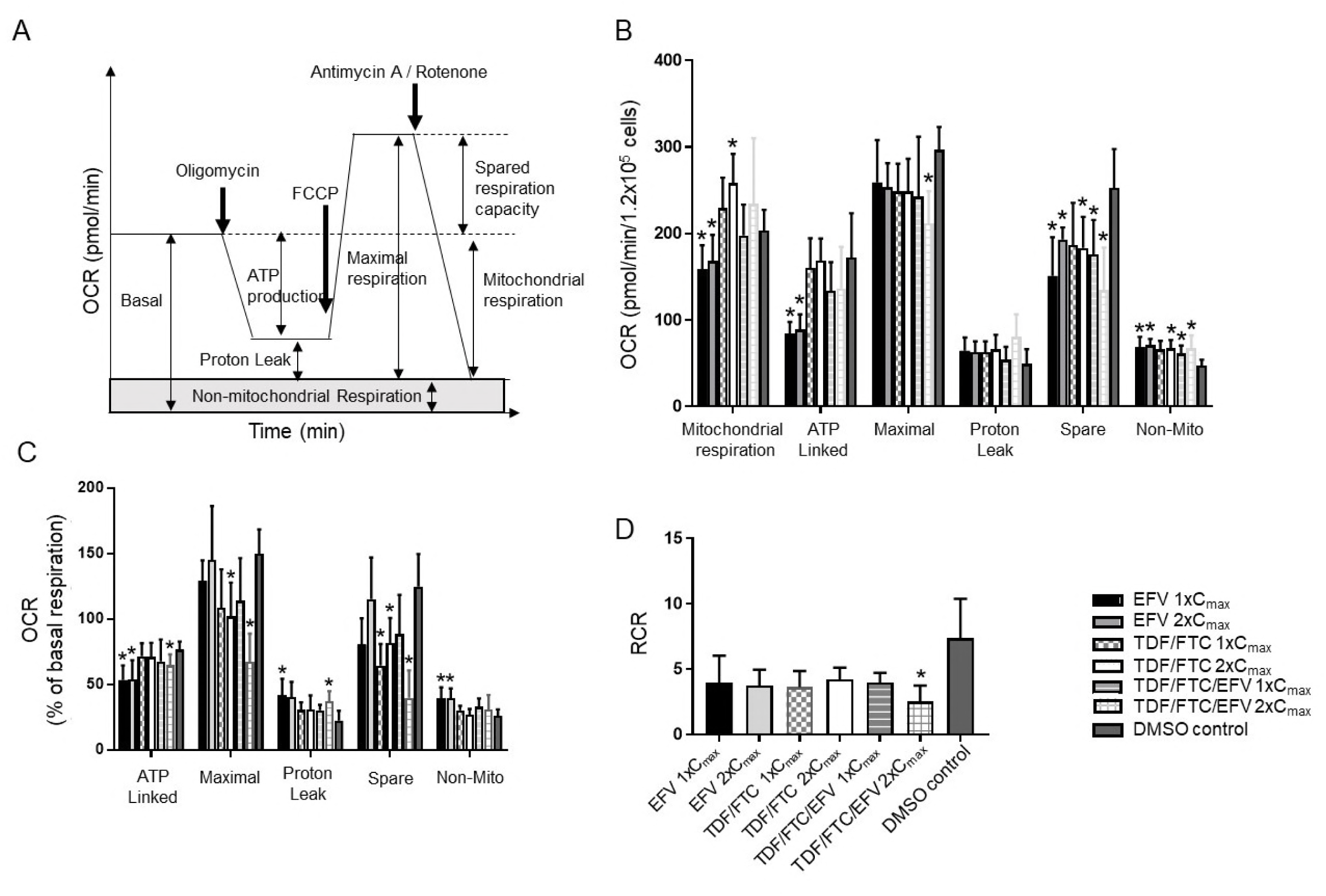
Effect of efavirenz treatment on mitochondrial respiratory function. Human T lymphoblastoid cell line (CEM cells) was treated with multiplicities of the plasma peak concentrations (1x-, or 2xC_max_) of Efaverinz (EFV), Tenofovir and Emtricitabine (TDF/FTC), and TDF/FTC/EFV for 2 days. The cells were counted and washed with the calibration medium and 1.2×10^5^ cell were added to each of 1% gelatin coated 96 wells in triplication. A sequential addition of specific inhibitors to mitochondria complex I, II, III and IV: 2 mM oligomycin (I), 0.6 mM FCCP (II), 0.4 mM FCCP (III) and 1 mM rotenone plus 1 mM antimycin A were done according to manufacturer’s instructions. A. Graphical representation of the OCR measurement over time. To estimate the proportion of OCR coupled to ATP synthesis, oligomycin (an inhibitor of ATP synthase—Complex V) was injected. To determine the maximal OCR, the proton ionophore (uncoupler) carbonylcyanide-4-(trifluoromethoxy)phenylhydrazone (FCCP) was injected, which stimulates respiration as the mitochondrial inner membrane becomes permeable to protons and electron transfer is no longer constrained by the proton gradient. Maximal OCR is controlled by ATP turnover (primarily involving the adenine nucleotide translocase, phosphate transporter and ATP synthase) and substrate oxidation–substrate uptake, processing enzymes, relevant ETC complexes, pool sizes of ubiquinone and cytochrome c, and O2 concentration. Finally, the high-affinity ETC inhibitors rotenone and antimycin A were added to inhibit the electron flux through Complex I and Complex III, respectively. The remaining OCR corresponds to the non-mitochondrial (or extramitochondrial) oxygen O_2_ consumption. B. Quantification of the mean OCR in CEM cells exposed to different treatment is shown for respiration under basal conditions (basal) and after the addition of oligomycin (proton leak) and FCCP (maximal respiration). The basal respiration rate minus the respiration rate recorded with oligomycin provides a measure of OCR due to ATP turnover whereas the respiration rate obtained with FCCP minus the basal O_2_ consumption values represents the reserve respiratory capacity. Non-mitochondrial respiration (rotenone plus antimycin A) was subtracted from each condition. C, Relative parameters of mitochondrial respiratory function and basal respiratory. The histograms show several parameters where data are expressed as the percentage of basal respiration (mitochondrial respiration plus non-mitochondrial respiration). D. Representation of the mean RCR. The RCR was calculated as the ratio between the maximal uncoupled mitochondrial respiration and mitochondrial respiration. Data (mean ±SD; n=3) were compared with DMSO treated cells and analyzed by Student’s t-test. P-values are two sided and considered significant if <0.05 (_*_), < 0.01 (_**_), or < 0.001 (_***_).

### Gene expression profiles of CEM cells treated with EFV, TDF, FTC, and TDF/FTC

We investigated the differential gene expression profiles of CEM cell treated with EFV (1x C_max_), TDF (4xC_max_), FTC (4xC_max_), and TDF/FTC (4xC_max_) using Affymetrix GeneChip Human Gene 2.0 ST arrays. We used 4xC_max_ of TDF and FTC because at lower concentrations, there was no significant differential gene expression. The number of genes expressed with at least 2-fold difference over cells treated with DMSO are illustrated in Figure 4A. There were no genes expressed at the 2-fold threshold with FTC treated cells. Total number of coding genes perturbed by TDF/FTC treatment was 3. EVF treatment led to perturbation of more coding genes (n=13); among these were 11 genes associated with lipid or cholesterol biosynthesis. Figures 4B and 4C illustrate the genes that were either down- or up-regulated by EFV and TDF/FTC treatment, respectively.

**Figure 4.**
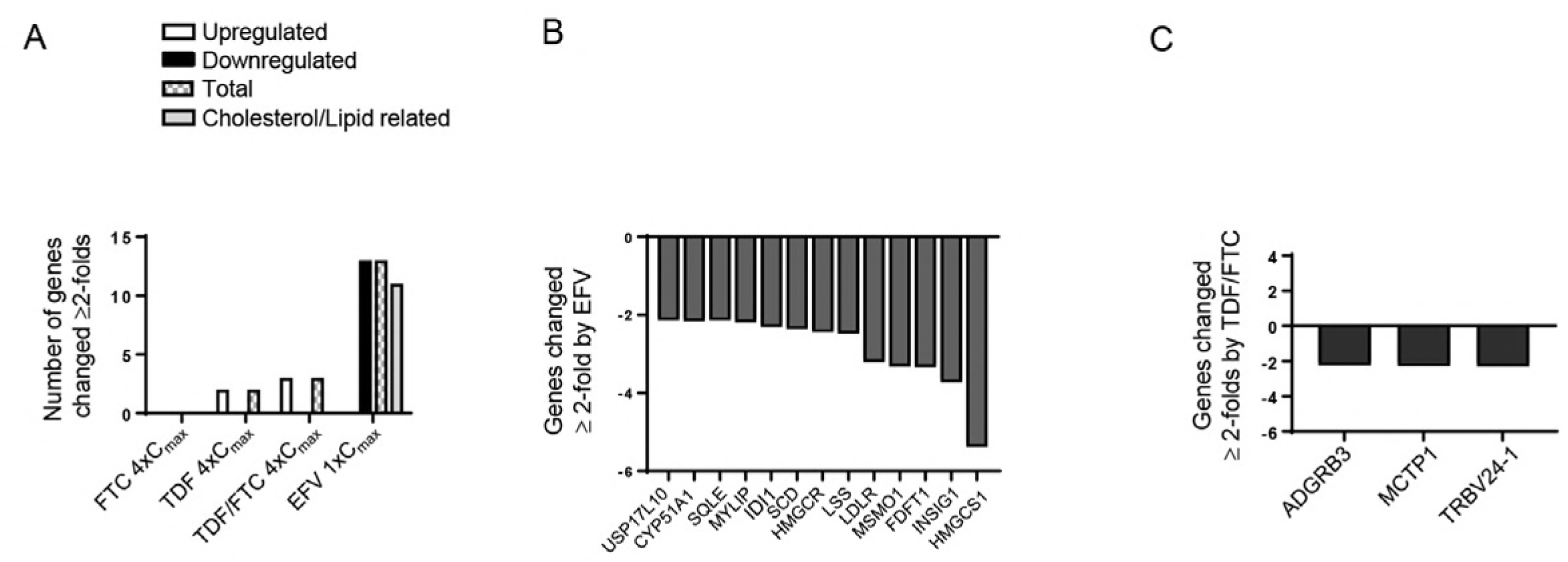
Effect of Efavirenz treatment on global gene expression. Human T lymphoblastoid cell line (CEM cells) was treated with 1xC_max_ of Efavirenz (EFV), 4xC_max_ of Tenofovir (TDF), or Emtricitabine (FTC) alone or in combination (TDF/FTC) for two days. The cells were collected and total RNA was extracted using TRIzol reagent. 100 ng of total RNA was amplified and labeled using the Whole-Transcript Sense Target Labeling Protocol. Affymetrix Gene Chip Human Gene 2.0 ST arrays were hybridized and scanned for signals. Each experiment was repeated on at least three occasions. Affymetrix Transcriptome Analysis Console TAC 3.1 was used to analyze the result. One-Way ANOVA was used and significance was achieved when fold change was ≥2 or ANOVA p-value < 0.05. A. The number of differentially expressed genes. B. Genes perturbed by treatment of CEM cells with EVF. C. Genes perturbed by treatment of CEM cells with TDF/FTC.

### EFV treatment was associated with down-regulation of genes associated with cholesterol biosynthesis

With the finding of downregulation of genes of cholesterol biosynthesis by EFV and previously reported effect of EFV on the lipogenic transcription factor sterol regulatory element binding protient-1c (SREBP-1c) (24), we sought to validate our microarray data with quantitative PCR (Figure 5). Relative gene expression in cells treated with antiretroviral drugs was compared to gene expression in cells treated with DMSO. Cells treated with EFV had a significant reduction in 3-hydroxy-3-methylglutaryl-CoA synthases, HMGCS, (p<0.01) and 3-hydroxy-3-methylglutaryl-CoA reductase, HMGCR, (p<0.05) expressions (Figure 5A). Combination treatment with TDF/FTV/EFV also decreased the expressions of HMGCS (p<0.01) and HMGCR (p<0.05). TDF/FTC treatment did not have any significant effect on the expressions of HMGCS and HMGCR. All treatment conditions had significant effect on insulin induced gene-1(INSIG-1) but not insulin induced gene-2 (INSIG-2) (Figure 5B). EFV treatment resulted in significant expressions of sterol regulatory element binding protien-1 (SREBP1) (p<0.05) (Figure 5C), low density lipoprotein receptor (LDLR) (p<0.05), and squalene epoxidase (SQLE) (p<0.05) (Figure 5D). None of the treatments compared to DMSO treatment had significant effect on the expressions of AMP-activated protein kinase (AMPK) –PRKAA1, PRKAA2, and PRKAB (Figure 5E).

**Figure 5.**
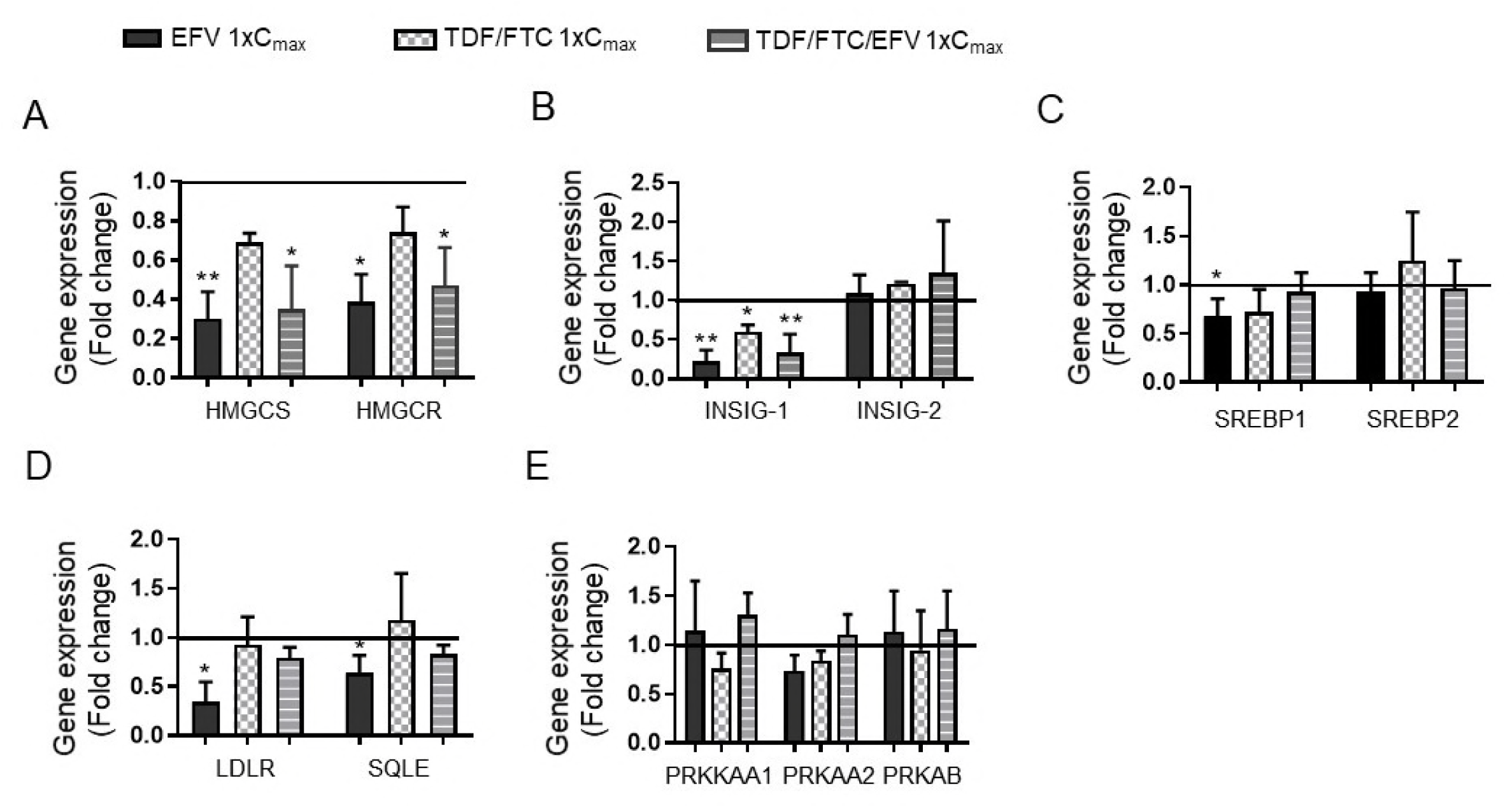
Effect of Efavirenz treatment on the expression of cholesterol biosynthesis. Human T lymphoblastoid cell line (CEM cells) was treated with multiplicities of the plasma peak concentrations (1x-, or 2xC_max_) of Efavirenz (EFV), Tenofovir and Emtricitabine (TDF/FTC), At day 2, the cells were harvested and RNA were extracted. Real-time RT-PCR of selected genes whose expression levels were normalized to the expression of housekeeping gene-GAPDH. Data (mean ± SD, n = 3), are represented as fold change in expression compared with cells treated with DMSO. A. Fold change in expressions of HMGCS and HMGCR; B. Fold change in expressions of INSIG-1 and INSIG-2; C. Fold change in expressions of SREBP1 and SREBP2; Fold change in expressions of LDLR and SQLE; and E. Fold change in expressions of PRKAA1, PRKAA2, and PRKAB. Data represent at least 3 independent experiments and plotted as mean ± SD. P-values are two sided and considered significant if <0.05 (_*_), < 0.01 (_**_), or < 0.001 (_***_).

## DISCUSSION

In this study, we sought to investigate whether combining EFV with TDF/FTC will moderate reported EFV-associated mitochondrial dysfunction (8, 18, 25-27) and perturbation of cholesterol biosynthesis (8, 28, 29). Most of the reported EFV-induced toxicities are from *in vitro* studies where cells were treated with EFV alone. However, in real life, EFV is used in combination with other ARVs such as TDF and FTC. Combination of TDF/FTC/EFV is one of the most popular ART regimens globally. We conducted comprehensive assessment of mitochondrial function from mtDNA content to mitochondrial respiration –using extracellular flux analysis, the gold standard for analyzing mitochondrial respiration in intact cells (18). EFV treated cells compared to DMSO treated cells had significant reduction in OCR contributed by mitochondrial respiration, ATP production-linked respiration, and SRC, increased apoptosis, high proportion of cells with reduced mitochondrial ΔΨ, increased production of ROS, decreased mtDNA content, and downregulation of genes involved in cholesterol biosynthesis in a dose- and time-dependent manner. Interestingly, treatment of cells with TDF/FTC/EFV combination had similar effect on mitochondrial function and cholesterol biosynthesis to treatment with EFV alone. Thus our findings do not support our hypothesis that agents given in combination with EFV moderate the effect of EFV on mitochondrial function leading to less prevalence of EFV-induced mitochondrial dysfunction in real time. If any, the effect of TDF and FTC on EFV-induced toxicity was marginal.

The threshold at which mitochondrial dysfunction will translate into toxic phenotype in a cell is not well understood; it depends on the type and energy requirement of the cell (30). In CEM cells treated with EFV, we observed a reduction in mitochondrial respiration, ATP production-linked respiration, and SRC (Figure 3B). These findings are consistent with effect of EFV on mitochondrial respiratory parameters in human glioma, neuroblastoma, and HepG2 cell lines (18, 31) and primary cultures of rat neurons and astrocytes (25) reported in previous studies. In these studies, EVF was cultured with the cells for 1 h to 24 h, while we extended treatment for 48 hrs. The SRC of a cell translates into the ability of the cell to survive and function under high energy demanding conditions (18). Decrease in SRC has been associated with heart diseases, neurodegenerative disorders and aging (32). Interestingly, the prevalence of these conditions is on the rise in HIV treatment-experienced individuals(33, 34). We also observed an increase in non-mitochondrial respiration with all treatment conditions (Figure 3B). Non-mitochondrial respiration accounts for other oxygen-consumption processes in the cell that do not involve the ETC. The increase in non-mitochondrial respiration suggests mitochondria dysfunction leading to a compromised ETC. Takemoto et al. recently reported that HIV-infected youth with insulin resistance had lower mitochondrial respiratory parameters and concluded that disordered mitochondrial respiration may be a potential mechanism of insulin resistant in HIV-infected youth (35).

Moreover, we investigated the effect of EFV on other functions of mitochondria. CEM cells treated with either EFV alone or in combination with TDF/FTC showed a higher proportion of non-viable cells (Figure 1A) and increased apoptosis (Figure 1B) in time- and dose-dependent manner. These findings are consistent with other reports; EFV has been reported to decrease cell viability and increase apoptosis in different cell types –primary human hepatocytes and HepG2 cell line (8, 36), pancreatic cancer cell lines (37), human endothelial cell lines (38), human T lymphocytes (Jurkat cell lines) (39), and PBMCs (39). Mitochondrial function depends on intact mitochondrial membrane (40). The loss of mitochondrial ΔΨ is believed to be the first and crucial step in mitochondrial dysfunction, which triggers a cascade of events leading ultimately to cell death(27, 41). It is, therefore, not surprising that the effect of EFV on mitochondrial ΔΨ (Figure 2A) paralleled the effect of EFV on apoptosis (Figure 1B). This is consistent with other studies that reported that treatment with EFV led to reduced mitochondrial ΔΨ in several cell types(25, 26). Mitochondrial dysfunction and compromise of ETC lead to increase production of ROS. Treat with EFV alone and in combination with TDF/FTC resulted in a time-dependent increase in ROS production. This finding is consistent with the effect of EFV on the production of ROS in hepatocytes (8). Treatment with EFV also resulted in about 50% reduction in mtDNA compared to treatment with DMSO. Although, EFV does not inhibit Pol-γ, we observed mtDNA depletion in CEM cells treated with 1xC_max_ of EFV; this is consistent with previous report using human hepatoma cells Huh 7.5 (27). Thus, EFV has several affections on mitochondria. In a study comparing Hep3B cells with functional mitochondria (rho+) and Hep3B cells lacking functional mitochondria (rho°), cells with functional mitochondria were more sensitive to the toxic effects of EFV (42).

Mitochondria play a role in cholesterol and lipid metabolism (43). In a microarray analysis, we observed a downregulation of the expression of cholesterol biosynthesis genes in CEM cells treated with EFV. We validated these data using qPCR assay to investigate the expression of genes involved in synthesis (HMGCS, HMGCR), regulation (SREBP1, SREBP2, INSIG-1, INSIG-2, PARKAA1, PRKAA2, PRKAB), and uptake (LDLR, SQLE). EFV treatment resulted in downregulation of HMGCS, HMGCR, INSIG-1, SREBP1, LDLR, and SQLE. Intracellular cholesterol is tightly regulated by cholesterol uptake, *de novo* synthesis, and efflux out of the cell (44). Our findings of downregulation of synthesis, uptake, and regulatory cholesterol biosynthesis genes imply that EFV might result in accumulation of intracellular cholesterol. These findings are consistent with the findings of Feeney et al. (44) in a case control study of expression of cholesterol biosynthesis genes in monocytes of HIV treatment-experienced, HIV treatment-naïve, and HIV-uninfected individuals. Blas-Garcia et al. also reported that EFV treatment of Hep3B cells and primary human hepatocytes resulted in intracellular accumulation of lipids (8). They concluded that EFV-induced mitochondrial dysfunction resulted in the activation of AMPK leading to intracellular accumulation of lipids. In our study, EFV did not have significant effect on AMPK (PRKAA1, PRKAA2, PRKAB) (Figure 5D). Moreover, we also observed that cells treated with EFV had decreased in the expression of INSIG-1. Under normal regulation of cholesterol homeostasis, depletion of intracellular cholesterol leads to increase expression of INSIG-1 (45). Our finding of decreased expression of INSIG-1, therefore, supports the hypothesis that EFV treatment may lead to intracellular accumulation of cholesterols. In contrast, Hadri et al. reported that treatment of human adipocytes with EFV resulted in depletion of intracellular triglycerides and at the same time a decrease in the expression of SREBP-1c (24). This differential effect of EFV on intracellular lipids may be due to the different types of cells or duration of treatment used in these studies. The C_max_ of EFV used in our experiments is about 12.7 µM, which is within the range of the plasma concentration of EFV in individuals receiving a daily dose of 600 mg of EFV (3.17 – 12.67 µM) (46). There is marked individual variability in plasma concentration of EFV; certain individuals can achieve plasma concentration over and above the 2xC_max_ used in our experiments. Individuals with certain polymorphisms in the cytochrome P450 (CP450) gene could have plasma concentrations ranging from 30 – 80 µM (47-50). The individual differences in EFV plasma concentration may partly explain why certain patients on EFV-based therapy do not experience effects of EFV on mitochondrial function. Ganta et al. detected EFV by HPLC in purified mitochondrial lysate from Huh 7.5 and HepaRG cells after 12 h of treatment with 6.25 µM (27). The investigators explained that since EFV is hydrophilic it can pass freely through the outer mitochondrial membrane and initiate depolarization of the inner mitochondrial membrane and subsequent mitochondrial dysfunction (27). This theory may be consistent with the individual differences in the mitochondrial affections of EFV.

Our study, like all *in vitro* studies, has several limitations. First, we could not possibly mimic or take into account all the complex biological pathways and processes that occur *in vivo*. Second, the experiments were conducted using a cancer cell line whose cellular bioenergetics may not be fully comparable to that of primary cells (18). We chose CEM cells because they can be infected with HIV. Therefore, they can be used to tease out the relative contributions of HIV infection and ART in mitochondrial dysfunction and cholesterol biosynthesis in future studies. Third, one might argue that 2 days of culture may not be adequate to tease out the delayed effect of EFV on mitochondria in real life. However, our duration is consistent with the duration used in several *in vitro* studies of ART toxicity (51, 52) and the decrease in viability of CEM cells with EFV treatment did not allow us to go beyond 48 h of culture.

In conclusion, EFV treatment resulted in reduction in mitochondrial respiratory parameters. Moreover, we observed increased apoptosis, high proportion of cells with reduced mitochondrial ΔΨ, increased production of ROS, decreased mtDNA content, and downregulation of genes involved in cholesterol biosynthesis in cells treated with EFV. Interestingly, combining TDF/FTC with EFV did not alter the effects of EFV on mitochondrial function. The main rationale of our study was to understand the gap between reported *in vitro* and *in vivo* EFV-induced toxicity. This gap may be due to individual differences in the pharmacokinetic of EFV. EFV can diffuse into the mitochondrial an initiate mitochondrial dysfunction (27). Thus, individuals who achieve higher tissue concentrations of EFV may have a predilection for EFV-induced mitochondrial dysfunction. Further studies are needed to elucidate the underlying mechanisms.

## METHODS AND MATERIALS

### Antiretroviral gents

EFV was obtained from the NIH AIDS Reagent Program (Germantown, MD, USA). FTC and TDF were purchased from Selleckchem (Houston, TX, USA). All the drugs were dissolved in dimethyl sulfoxide (DMSO). DMSO was purchased from Sigma-Aldrich (St. Louis, Missouri, USA).

### Cell culture and cell viability

Human T lymphoblastoid cell line (CEM cells) was cultured in RPMI 1640 supplemented with 10% dialyzed fetal calf serum (Thermo Fisher Scientific, NY, USA). Cells were incubated at 37ºC in a 5% CO_2_ humidified environment. 3 x 10^4^/ml cells (total volume of 40 ml) were incubated with EFV, FTC, or TDF at multiples of plasma peak concentration, e.g., 1xC_max_, or 2xC_max_. The cells were collected for cell count and the experiments described below each day. Aliquots of cells were stained with Trypan blue to distinguish live cell from dead cells and the number of live cells was counted using hemocytometer. For each treatment condition at least three cell culture experiments were conducted on different occasions.

### Apoptosis assay

Apoptosis was assessed by staining with an FITC Annexin V Apoptosis Detection Kit I (BD, San Jose, CA) according to the manufacturer’s instructions. In brief, the cells cultured with various concentrations of EFV, TDF/FTC, or TDF/FTC/EFV (1x-, or 2x-C_max_) were harvested at day 1 and 2 and then 1 × 10^6^ cells were washed with ice-cold PBS. The early apoptotic cells (Annexin V+/PI-) or late apoptotic cells (Annexin V+/PI+) were evaluated by double staining with annexin V–FITC and PI and then run on BD LSR II flow cytometer and analyzed with BD FACSDiva software (San Jose, CA). Each experiment was done in triplicates and repeated on at least three occasions.

### Mitochondrial membrane potential (ΔΨ)

Mitochondrial function depends on intact mitochondria membranes that can maintain an electrochemical potential generated by the electron transport chain, therefore, mitochondrial ΔΨ is a good measure of mitochondrial functional capacity. Mitochondrial ΔΨ was analyzed using TMRE-Mitochondrial Membrane Potential Assay Kit (Abcam, Cambridge, MA) according to the manufacturer’s instructions. TMRE (tetramethylrhodamine, ethyl ester) is a cell permeant, positively-charged, red-orange dye that readily accumulates in active mitochondria due to their relative negative charge. Depolarized or inactive mitochondria have decreased membrane potential and fail to sequester TMRE. At day 1 and 2 of treatment, cells were collected and stained with 200 nM TMRE respectively for 20 min at room temperature. The reaction was stopped by adding 300 µl of PBS and immediately analyzed on BD LSR II flow cytometer (BD, San Jose, CA) and with BD FACS Diva software. Each experiment was done in triplicates and repeated on at least three occasions.

### Mitochondrial reactive oxygen species production

Production of ROS by mitochondria was determined using MitoSOX™ Red mitochondrial superoxide indicator, a novel fluorogenic dye for highly selective detection of superoxide in the mitochondria of live cells, accordingly to manufacturer’s instructions. Treated cells were washed once with wash buffer before incubation with 5 μM MitoSOX™ reagent working solution at room temperature for 10 min. The cells were then washed and run on LSRII flow cytometry (Beckman Coulter). Each experiment was done in triplicates and repeated on at least three occasions.

### Mitochondrial DNA quantification

To assess mtDNA content, genomic DNA was extracted from CEM cells treated with EFV, TDF/FTC, or TDF/FTC/EFV using TRIzol® Reagent according to manufacturer’s instructions. Fragment of MT-TL1gene (encodes tRNA leucine 1) and 18 S rRNA nuclear gene were amplified using quantitative RT-PCR as previously described (53). The primers for mitochondrial fragment (MIT) and 18S were (MIT Forward, 5’ AGGACAAGAGAAATAAGGCC and MIT Reverse, 5’ TAAGAAGAGGAATTGAACCTCTGACTGTAA) and (18S Forward, 5’ TAC CTG GTT GAT CCT GCC AGT and 18S Reverse, 5’ GAT CCT TCC GCA GGT TCA CCT AC), respectively. Each experiment was done in duplicate and repeated on at least three occasions. The relative mtDNA content was determined as the ratio of mtDNA to nuclear DNA. Final data for each treatment condition represent the mean and standard deviation (SD) from at least three cell culture experiments.

### Mitochondrial respiratory function

Mitochondrial respiratory function was determined using the Seahorse XF-96 Extracellular Flux Analyzer (Agilent Technologies, Inc., Wilmington, DE, USA) according to manufacturer’s instructions (www.agilent.com/en-us/products/cell-analysis-(seahorse)/basic-procedures-to-run-an-xf-assay) and as previously published (18, 35). In brief, the Agilent Seahorse uses modulators of respiration that target components of the electron transport chain. The compounds (oligomycin, FCCP, and a mix of rotenone and antimycin A) were serially injected to measure ATP production, maximal respiration, and non-mitochondrial respiration, respectively. Proton leak and spare respiration capacity (SRC) were then calculated using these parameters and basal respiration. Treated CEM cells were seeded at a density of 1.2 x 10^5^ cells per well in triplicate in a 1% gelatin treated 96 -well plate. Then, the sensor cartridge was load with oligomycin, FCCP, and a mixture of rotenone and antimycin A in ports A, B, and C, respectively. The cell culture plate was then inserted into the analyzer and programed to ensure a homogenous environment. For each phase, measurements were performed in triplicate, totaling 12 points per trace. To allow for comparison between different experiments, the data obtained for each condition were normalized to the cell number per well and expressed as the oxygen consumption rate (OCR) in pmol/min. The experiment was repeated three times. No significant inter-assay differences were detected between the three repeats with regard to appearance and viability of cell or the values of OCR that were obtained.

### Microarray gene expression assay

Total RNA was extracted from cell pellets using TRIzol (Life Technologies, Rockville, MD) according to manufacturer’s instructions. Contaminated genomic DNA was removed by treating RNA samples with Ambion RNase-free DNase (Thermo Fisher Scientific) for 20 min at 37 C. RNA quantity and quality were assessed by micro-volume spectrophotometry on an Infinite 200 PRO plate reader (Tecan, Männedorf, Switzerland) and by on-chip capillary electrophoresis on a Bioanalyzer 2100 (Agilent Technologies, Santa Clara, CA), respectively. Absorbance ratio at 260 and 280 nm was ≥1.9 and the RNA integrity number (RIN) was >8 for all the samples. 100 ng of total RNA was amplified and labeled using the Whole-Transcript Sense Target Labeling Protocol by Affymetrix (Santa Clara, CA) without ribosomal RNA reduction. Affymetrix GeneChip Human Gene 2.0 ST arrays were hybridized with 11 µg of labeled sense DNA, washed, stained, and scanned on an Affymetrix 7G Scanner according to the manufacturer’s protocols. Each experiment was repeated on at least three occasions. Affymetrix Transcriptome Analysis Console TAC 3.1 was used to analyze the result. One-Way ANOVA was used and significance was achieved when fold change was ≥2 and a p-value of < 0.05.

### Quantitative real-time polymerase chain reaction for expression of cholesterol biosynthesis genes

RNA was extracted from aliquots of EFV-, TDF/FTC-, or TDF/FTC/EFV-treated cells using TRIzol Reagent (ThermoFisher Scientific, Carlsbad, CA) according to the manufacturer’s instructions and used in quantitative real-time PCR as previously described (54). Melting curve analysis was performed after the completion of PCR to assess the possibility of false-positive results. All of the samples were run in duplicate in at least three independent experiments. The genes of interest were HMGCS, HMGCR, SREBP-1 and -2, INSIG-1 and -2, LDLR, SQLE) and AMPK – PRKAA1, PRKAA2 and PRKAB. The primers are listed in Supplementary Table 1. The gene encoding glyceraldehyde 3-phosphate dehydrogenase (GAPDH) was used as an internal control for all reactions. The threshold cycle (CT) values of the genes were determined for each treatment condition. The fold-change in gene expression was calculated as 2^∆∆CT^; where ∆∆CT = ∆CT_(treated)_ – ∆CT_(control)_; ∆CT_(treated)_ = (CT_(gene of interest)_ – CT_(GAPDH)_); ∆CT_(control)_= (CT_(gene of interest)_ – CT_(GAPDH)_).

### Data and statistical analysis

All statistical analyses were performed with GraphPad Prism software with the Student’s t-test. Data are expressed as means ± SD and significance was achieved when p value was <0.05.

## Acknowledgements

This study was supported by grants from the National Institutes of Health (KO8AI074404 and R01 HD074252 to EP). AS was supported by an IDSA Medical Scholar Award.

